# Multifactorial pathways facilitate resilience among kindergarteners at risk for dyslexia: A longitudinal behavioral and neuroimaging study

**DOI:** 10.1101/618298

**Authors:** Jennifer Zuk, Jade Dunstan, Elizabeth Norton, Xi Yu, Ola Ozernov-Palchik, Yingying Wang, Tiffany P. Hogan, John D. E. Gabrieli, Nadine Gaab

## Abstract

Recent efforts have focused on screening methods to identify children at risk for dyslexia as early as preschool/kindergarten. Unfortunately, while low sensitivity leads to under-identification of at-risk children, low specificity can lead to over-identification, resulting in inaccurate allocation of limited educational resources. The present study focused on children identified as at-risk in kindergarten who do *not* subsequently develop dyslexia to specify factors associated with better reading outcomes among at-risk children. Early screening was conducted in kindergarten and a subset of children was tracked longitudinally until second grade. Potential protective factors were evaluated at cognitive-linguistic, environmental, and neural levels. Relative to at-risk kindergarteners who subsequently developed dyslexia, those who did not were characterized by significantly higher socioeconomic status (SES), speech production accuracy, and microstructure of the posterior right-hemispheric superior longitudinal fasciculus (SLF). A positive association between microstructure of the right SLF and subsequent decoding skills was found to be specific to at-risk children and not observed among typical controls. Among at-risk children, several kindergarten-age factors were found to significantly contribute to the prediction of subsequent decoding skills: microstructure of the posterior right SLF, age, gender, SES, and phonological awareness. These findings suggest that putative compensatory mechanisms are already present by the start of kindergarten. The right SLF, in conjunction with the cognitive-linguistic and socioeconomic factors identified, may play an important role in facilitating reading development among at-risk children. This study has important implications for approaches to early screening, and assessment strategies for at-risk children.

## INTRODUCTION

### The importance of early identification of children at risk for dyslexia

Significant disparities in early reading achievement prevail in the United States, as nearly two thirds of fourth grade students demonstrate reading skills below grade level^1^. Children who significantly struggle with decoding, speed, and accuracy of single word reading may be characterized by developmental dyslexia (henceforth referred to as dyslexia), a specific learning disorder that cannot be explained by sensory, motor or cognitive deficits, lack of motivation, or deficient educational opportunities^2,3^. Presently, dyslexia is often only diagnosed after a child demonstrates significant difficulty learning to read, and as such, diagnosis typically occurs in late elementary school^4^. However, persistent discouraging experiences with reading can have negative psychosocial consequences such as feelings of frustration and helplessness, as well as higher rates of depression and anxiety^5^, which can impact academic achievement long-term and hinder vocational potential^6^.

Early screening at the start of formal reading instruction (i.e., kindergarten classrooms in the United States) or even earlier offers great potential to identify at-risk children and facilitate reading development through early intervention^7^. One of the most reliable markers of dyslexia prior to reading onset is poor phonological awareness, the ability to manipulate speech sounds within oral language^8^. In addition, key predictors of subsequent literacy outcomes in early childhood include letter-sound knowledge and rapid automatized naming skills^9–16^. Moreover, additional factors may contribute to difficulty learning to read, such as speech and receptive and expressive language delays^12,16–20^. Approximately 50-90% of children who are identified early as at-risk and provided with targeted instruction have been shown to achieve grade-level reading skills thereafter^21–24^. Although these promising benefits have been observed, current approaches to early identification often tend to identify too many children as being ‘at-risk,’ meaning that many ‘at-risk’ children as identified by early screening do not end up developing dyslexia^25–27^, which can lead to inaccurate allocation of educational resources. This raises an important question of who is *truly* at risk. Among children who are initially identified as at-risk based on early screening, what factors distinguish those who do *not* go on to develop dyslexia relative to those who do? The present study focused on characterizing at-risk children who do not subsequently develop dyslexia to examine the factors that are associated with better word reading outcomes among children who have been classified as being at-risk based on screening protocols.

### The dynamic, multi-factorial trajectory of dyslexia

Converging evidence supports the notion of a liability distribution for dyslexia that is determined by an interaction between multiple factors at cognitive, neural, genetic, and environmental levels^28,29^. It has been suggested that a probabilistic interaction of several risk factors increases the liability to develop dyslexia; meanwhile, proposed ‘protective’ factors may decrease the liability^28,29^. Consequently, not all children identified as ‘at-risk’ early on subsequently develop dyslexia due to the dynamic interaction of these factors, which ultimately give rise to variable outcomes ranging from good to poor reading skills^16^. Although this holds great promise for facilitating reading development among at-risk children, it remains unclear what specific factors may be associated with better reading outcomes. Delineation of these factors offers the potential to guide educational and clinical approaches to effectively identify at-risk children and subsequently support their reading development.

### Factors that may promote successful reading acquisition among at-risk children

#### Cognitive-linguistic factors

In addition to the factors shown to be most predictive of general reading outcome, several cognitive-linguistic factors have been proposed to promote reading development among children at risk for dyslexia^16,30^. Children with familial risk for dyslexia who go on to develop typical reading skills relative to those who develop dyslexia have shown enhanced cognitive and oral language abilities, particularly in terms of vocabulary knowledge^31–36^ and syntactic structure^32,34,35^. Relative strengths in these components of language may further support phonological development, and/or be utilized as a strategy to achieve word recognition during reading acquisition^16,37^.

Speech-sound production accuracy has also been put forth as a potential factor, though investigation to date has only shown trends towards significance^31,32,35,36^. Although significant differences have yet to be observed, it has been hypothesized that the currently available standard articulation tests may not be sensitive enough to discern group differences in speech production accuracy between those who go on to have dyslexia and those who do not^31,32^.

#### Environmental factors

Certain environmental and family-related factors have also been strongly linked with reading development, particularly those related to socioeconomic status^38–40^. Research has repeatedly shown the critical impact of socioeconomic status on language and reading development^38,41,42^. From early childhood, significant disparities in vocabulary knowledge have been observed when comparing children from high versus low socioeconomic backgrounds^43,44^. This achievement gap has been documented to widen along the developmental trajectory, presumably due to the complex influence of parent education level, occupation, and income on child development^45–47^.

Ultimately, reading acquisition is one of many academic skills impacted by socioeconomic factors, and higher socioeconomic status is positively linked with language (especially vocabulary knowledge) and reading development^48–50^. However, many of the longitudinal studies tracking reading development among at-risk children have either recruited a sample that lacks a broad socioeconomic distribution, or have established comparison groups that do not differ in socioeconomic status^16^. Therefore, investigation with at-risk children from a broad socioeconomic sample is needed to further delineate the extent to which disparities in socioeconomic status play a significant role among at-risk children in shaping their subsequent reading outcomes.

#### Putative right-hemispheric neural pathways that facilitate reading development in at-risk children and those with dyslexia

In addition to cognitive-linguistic and environmental factors, right-hemispheric neural pathways have also been proposed to underlie better reading outcomes among at-risk children^51^. White matter has been intricately linked with reading development, as measured by diffusion-weighted imaging^52–54^. Several left-hemispheric pathways have been shown to significantly differ among children and adults with dyslexia compared to their peers with typical reading skills in terms of fractional anisotropy FA, the degree of directionality of water diffusion within white matter tracts^52,55,56^.

Although predominantly left-hemispheric pathways have been identified in comparisons of typical readers to those with dyslexia, focusing solely on the reading outcomes of children with dyslexia revealed that FA in the *right-hemispheric* superior longitudinal fasciculus significantly predicted reading improvements after approximately two years of reading instruction^57^. This relationship was not present among typical readers, thereby specific to children with dyslexia. Thus, investigation with a focus on the neural substrates linked with reading improvements among children with dyslexia revealed that right-hemispheric pathways seem to play an important role in facilitating better reading outcomes. Yet, it remains unclear whether pathways develop over the course of learning to read to compensate for difficulties, or whether they may be evident from the start of reading onset and predict reading outcomes among at-risk children.

Investigating white matter structure among at-risk children from the start of formal reading instruction remains understudied, yet offers great potential to better understand the pathways that underlie typical reading development. Group comparison between children with familial risk for dyslexia and those without has revealed significant differences in left-hemispheric white matter pathways even before children start learning to read^58–61^. However, scarcely any studies have focused primarily on at-risk children to examine how white matter microstructure at the start of formal reading instruction may be associated with subsequent reading development, particularly for at-risk children who do *not* go on to develop dyslexia.

Of the limited evidence to date, preschool age children with familial risk for dyslexia who subsequently became good readers were found to show a significantly higher developmental rate in the right-hemispheric superior longitudinal fasciculus compared to those who became poor readers^59^. This initial evidence suggests that among children with familial risk for dyslexia, right-hemispheric white matter pathways in particular are positively linked with better reading outcomes. Yet, it remains unclear to what extent this may be evident among at-risk children identified based on behavioral predictors of dyslexia (i.e., early screening), and whether this pathway may show altered microstructure from the start of reading instruction. Moreover, no study to date has investigated microstructure of this pathway concurrently with factors on cognitive-linguistic and environmental levels.

### Purpose of the present study

Several factors have been suggested to facilitate successful reading development among children at risk for dyslexia on cognitive-linguistic, environmental, and neural levels; however, there has yet to be a study to examine these factors concurrently or within the context of classifying risk status based on early screening. Therefore, the present study conducted screening in kindergarten classrooms in New England in a variety of schools with broad socioeconomic representation, and then enrolled a carefully selected subset of these children in a longitudinal investigation until the end of second grade. At-risk children were identified in kindergarten based on performance on key behavioral predictors of dyslexia (i.e., phonological awareness, rapid automatized naming, letter-sound knowledge^9–16^. Neuroimaging was acquired at the kindergarten age, and then children were longitudinally tracked and assessed with a comprehensive battery of standardized behavioral assessments. This study design allowed for investigation of cognitive-linguistic, environmental, and neural factors associated with subsequent word reading abilities among at-risk children identified at the start of formal reading instruction.

To address important missing links in the literature, the multidimensional approach employed in the present study investigated a few sequential research questions. The first research question of interest was: among at-risk children identified based on early screening in kindergarten, what factors differ between at-risk children who subsequently develop dyslexia, relative to those who do not? It is hypothesized that putative factors from previous research, particularly in cognitive-linguistic domains^16^, will be reflected in the present analysis focused on a community sample with determination of risk status conducted in a manner that may align with educational practices. Among these factors, speech production accuracy is included with the implementation of an alternative to standard articulation^31,32^, utilizing detailed analysis of speech production accuracy through audio-recorded speech samples. In addition, rather than matching groups based on socioeconomic status, we hypothesized that socioeconomic status would significantly differ between at-risk children who subsequently develop dyslexia relative to those who do not.

The second component of the present study focused on the previously hypothesized role of right-hemispheric superior longitudinal fasciculus in facilitating better reading outcomes among children at risk/with dyslexia. To build upon previous evidence, the present study asked the following questions: what is the relationship between microstructure in the right superior longitudinal fasciculus at the kindergarten age and subsequent word reading abilities in second grade? Are these relationships specific to at-risk children, or are they evident among typical controls as well? Based on previous studies in older children and children with a familial risk, positive significant relationships are hypothesized only among at-risk children specifically^57,59^. Lastly, the final component of this study examined behavioral and neural factors concurrently to evaluate which variables at the kindergarten age best predict subsequent word reading abilities among at-risk children. Taken together, this work has the potential to identify protective factors that may underlie the trajectory of at-risk children who do *not* subsequently develop dyslexia, and reveal which variables on cognitive-linguistic, environmental, and neural levels best predict subsequent word reading outcomes among at-risk children. In turn, this study will inform educational and clinical approaches to early identification, as well as supporting reading development in at-risk children from the start of formal reading instruction.

## RESULTS

### Among at-risk children, how do children who subsequently do not develop dyslexia differ from those who do, relative to typical controls?

#### Initial assessment at the start of formal reading instruction

A one-way MANOVA was employed to compare performance on all kindergarten-age factors between at-risk children with subsequent dyslexia, at-risk children without dyslexia, and typical controls. Regarding classification variables utilized for early screening, as expected, the one-way MANOVA verified group differences that reflect our approach to group categorization. Specifically, FDR-corrected post-hoc tests confirmed that typical controls performed significantly better than both at-risk groups on measures of phonological awareness, rapid automatized naming, and letter-sound knowledge (*p* < 0.05; summary of MANOVA provided in Table 1), and at-risk groups did not significantly differ on any of these screening measures. Word identification abilities in kindergarten showed a similar pattern of group differences, such that typical controls demonstrated significantly better abilities than both at-risk groups (*p* < 0.05), and at-risk children without subsequent dyslexia did not significantly differ in word identification from those who went on to develop dyslexia (*p* > 0.1). Notably, a similar gender distribution was observed among all three groups.

**Table 1.**
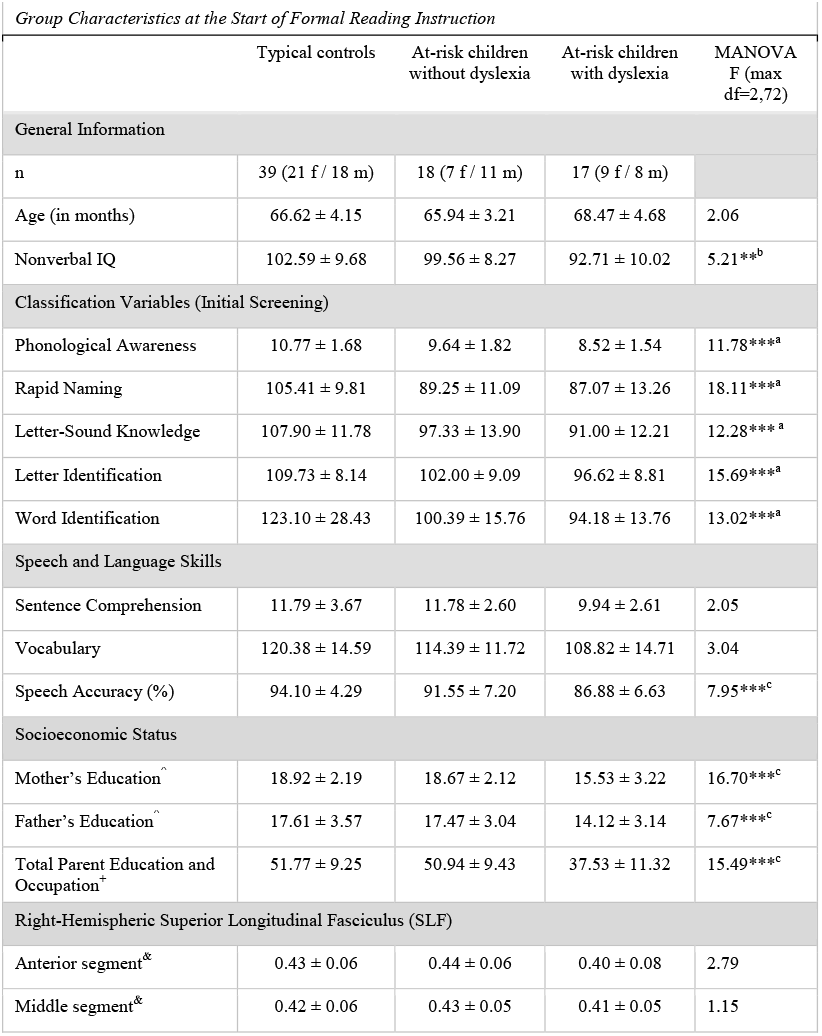
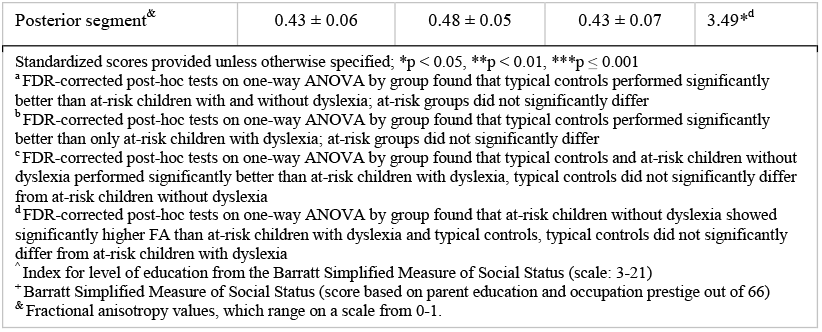
Group Characteristics at the Start of Formal Reading Instruction

|  | Typical controls | At-risk children without dyslexia | At-risk children with dyslexia | MANOVA<br>F (max<br>df=2,72) |
| --- | --- | --- | --- | --- |
| General Information |  |  |  |  |
| n | 39 (21 f / 18 m) | 18 (7 f / 11 m) | 17 (9 f / 8 m) |  |
| Age (in months) | 66.62 ± 4.15 | 65.94 ± 3.21 | 68.47 ± 4.68 | 2.06 |
| Nonverbal IQ | 102.59 ± 9.68 | 99.56 ± 8.27 | 92.71 ± 10.02 | 5.21**<sup>b</sup> |
| Classification Variables (Initial Screening) |  |  |  |  |
| Phonological Awareness | 10.77 ± 1.68 | 9.64 ± 1.82 | 8.52 ± 1.54 | 11.78***<sup>a</sup> |
| Rapid Naming | 105.41 ± 9.81 | 89.25 ± 11.09 | 87.07 ± 13.26 | 18.11***<sup>a</sup> |
| Letter-Sound Knowledge | 107.90 ± 11.78 | 97.33 ± 13.90 | 91.00 ± 12.21 | 12.28***<sup>a</sup> |
| Letter Identification | 109.73 ± 8.14 | 102.00 ± 9.09 | 96.62 ± 8.81 | 15.69***<sup>a</sup> |
| Word Identification | 123.10 ± 28.43 | 100.39 ± 15.76 | 94.18 ± 13.76 | 13.02***<sup>a</sup> |
| Speech and Language Skills |  |  |  |  |
| Sentence Comprehension | 11.79 ± 3.67 | 11.78 ± 2.60 | 9.94 ± 2.61 | 2.05 |
| Vocabulary | 120.38 ± 14.59 | 114.39 ± 11.72 | 108.82 ± 14.71 | 3.04 |
| Speech Accuracy (%) | 94.10 ± 4.29 | 91.55 ± 7.20 | 86.88 ± 6.63 | 7.95***<sup>c</sup> |
| Socioeconomic Status |  |  |  |  |
| Mother's Education<sup>^</sup> | 18.92 ± 2.19 | 18.67 ± 2.12 | 15.53 ± 3.22 | 16.70***<sup>c</sup> |
| Father's Education<sup>^</sup> | 17.61 ± 3.57 | 17.47 ± 3.04 | 14.12 ± 3.14 | 7.67***<sup>c</sup> |
| Total Parent Education and Occupation<sup>+</sup> | 51.77 ± 9.25 | 50.94 ± 9.43 | 37.53 ± 11.32 | 15.49***<sup>c</sup> |
| Right-Hemispheric Superior Longitudinal Fasciculus (SLF) |  |  |  |  |
| Anterior segment<sup>&</sup> | 0.43 ± 0.06 | 0.44 ± 0.06 | 0.40 ± 0.08 | 2.79 |
| Middle segment<sup>&</sup> | 0.42 ± 0.06 | 0.43 ± 0.05 | 0.41 ± 0.05 | 1.15 |
| Posterior segment<sup>&</sup> | 0.43 ± 0.06 | 0.48 ± 0.05 | 0.43 ± 0.07 | 3.49*<sup>d</sup> |

Standardized scores provided unless otherwise specified; \*p < 0.05, \*\*p < 0.01, \*\*\*p ≤ 0.001
<sup>a</sup> FDR-corrected post-hoc tests on one-way ANOVA by group found that typical controls performed significantly better than at-risk children with and without dyslexia; at-risk groups did not significantly differ
<sup>b</sup> FDR-corrected post-hoc tests on one-way ANOVA by group found that typical controls performed significantly better than only at-risk children with dyslexia; at-risk groups did not significantly differ
<sup>c</sup> FDR-corrected post-hoc tests on one-way ANOVA by group found that typical controls and at-risk children without dyslexia performed significantly better than at-risk children with dyslexia, typical controls did not significantly differ from at-risk children without dyslexia
<sup>d</sup> FDR-corrected post-hoc tests on one-way ANOVA by group found that at-risk children without dyslexia showed significantly higher FA than at-risk children with dyslexia and typical controls, typical controls did not significantly differ from at-risk children with dyslexia
<sup>^</sup> Index for level of education from the Barratt Simplified Measure of Social Status (scale: 3-21)
<sup>+</sup> Barratt Simplified Measure of Social Status (score based on parent education and occupation prestige out of 66)
<sup>&</sup> Fractional anisotropy values, which range on a scale from 0-1.

Investigation of potential protective or compensatory factors revealed group differences on cognitive-linguistic, environmental, and neural levels (for an overview see Table 1), as follows:

*On the cognitive-linguistic level*, significant effects in the MANOVA were observed for nonverbal cognitive abilities (F(2,72) = 5.21; *p* < 0.01) and speech production accuracy (F(2,72) = 7.95; *p* = 0.001). In terms of nonverbal cognitive abilities, FDR-corrected post-hoc tests revealed significantly better performance in typical controls relative to at-risk children with dyslexia (*p* < 0.005) and no differences between at-risk groups. Yet, post-hoc comparisons for speech production accuracy revealed significant differences between the at-risk groups, such that at-risk children without dyslexia did not differ from typical controls, and both groups showed significantly better accuracy relative to at-risk children with subsequent dyslexia (*p* < 0.05). By contrast, no significant group differences were observed in terms of vocabulary knowledge or sentence comprehension at the kindergarten age.

*On the environmental level*, significant group differences were also observed in all aspects of socioeconomic status: mother’s education level (F(2,72) = 16.70; *p* < 0.001), father’s education level (F(2,72) = 7.67; *p* = 0.001), and the index indicating total parent education and occupation prestige levels (F(2,72) = 15.49; *p* < 0.001). FDR-corrected post-hoc tests revealed that at-risk children without subsequent dyslexia did not differ from typical controls (*p* > 0.5), and both of these groups showed significantly higher socioeconomic levels in all aspects relative to at-risk children who went on to develop dyslexia (*p* ≤ 0.005).

*On the neural level*, fractional anisotropy (FA) in the posterior segment of the right superior longitudinal fasciculus (SLF) revealed significant group differences (F(2,72) = 3.49; *p* < 0.05; see Figure 1). Interestingly, at-risk children who did not subsequently develop dyslexia were characterized by significantly greater FA in this portion of the tract when compared to both typical controls and at-risk children who went on to develop dyslexia (*p* < 0.05), whereas typical controls did not differ from at-risk children with dyslexia (*p* > 0.5). Table 1 provides a summary of all group comparisons at the kindergarten age.

**Figure 1 Legend:**
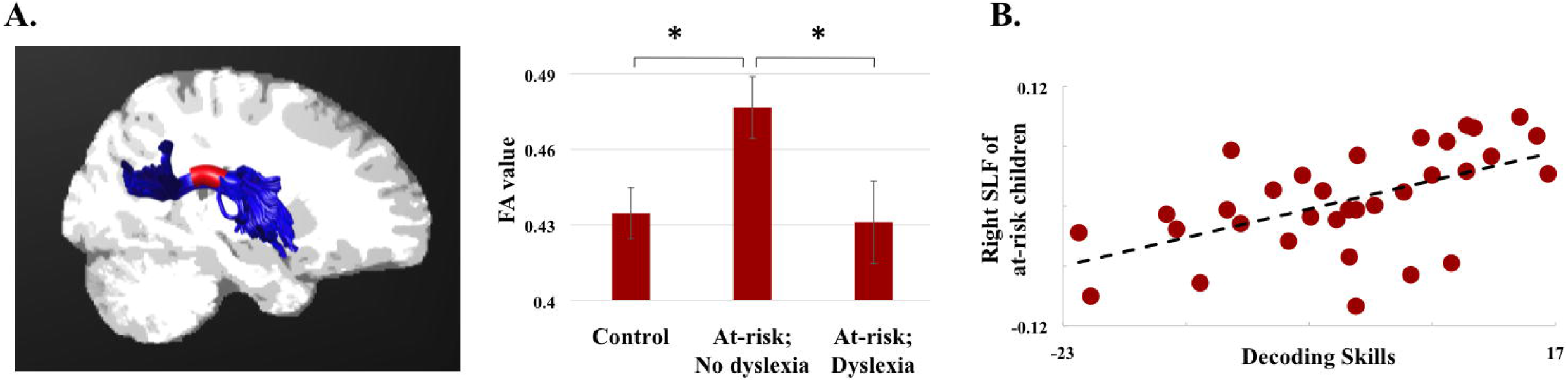
A. Significant group differences in the posterior segment of the right superior longitudinal fasciculus (SLF), signified in red in display. B. Significant partial correlation among all at-risk children between FA of the posterior segment of the right-hemispheric SLF in kindergarten and decoding skills at the end of second grade (displayed in terms of the centered residuals produced from partial correlations).

#### Longitudinal follow-up at the end of second grade

At the end of second grade, a MANOVA was employed with reading measures to verify group classification, and evaluate respective language abilities upon longitudinal follow-up. As expected based on our approach to group classification of subsequent reading outcomes, at-risk children without subsequent dyslexia and typical controls performed significantly better than at-risk children with dyslexia on all timed and untimed reading measures (FDR-corrected post-hoc tests, *p* < 0.001), and at-risk children who did *not* develop dyslexia did not significantly differ from typical controls on any of these measures.

In terms of general language abilities at the end of second grade, significant group differences reflective of reading outcomes were observed. The MANOVA revealed significant differences in sentence comprehension (F(2,72) = 7.55; *p* = 0.001) and vocabulary knowledge (F(2,72) = 9.88; *p* < 0. 001). Specifically, FDR-corrected post-hoc tests revealed that typical controls and at-risk children who did *not* develop dyslexia did not differ from each other on either measure, but the significant differences between at-risk groups varied by measure. For sentence comprehension, at-risk children with versus without subsequent dyslexia did not significantly differ, though there was a trend towards better performance among at-risk children who did *not* develop dyslexia (*p* < 0.1). Accordingly, in terms of vocabulary knowledge, at-risk children without subsequent dyslexia demonstrated significantly higher performance than at-risk children with dyslexia (*p* < 0.01). For a full summary of group comparisons at this time point, see Table 2.

**Table 2.**
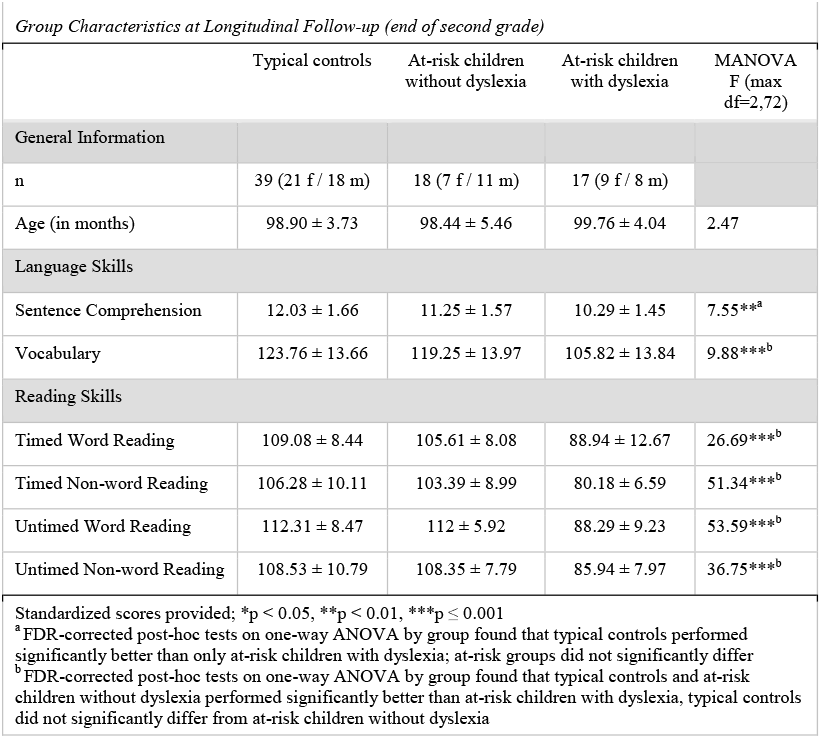
Group Characteristics at Longitudinal Follow-up (end of second grade)

### Does FA within the right SLF in kindergarten significantly relate to subsequent word reading abilities at the end of second grade among at-risk children?

To examine relationships between the right SLF in kindergarten and subsequent word reading measures in second grade, partial correlations between microstructure of the right-hemispheric SLF (as indicated by fractional anisotropy, FA) in kindergarten and subsequent timed and untimed word/non-word reading scores were employed among at-risk children only. Partial correlations were employed in order to examine these relationships independent of the effects of age, nonverbal cognitive abilities, and socioeconomic status. To further confirm whether relationships were specific to at-risk children only, these partial correlations were employed among typical controls as well.

Among typical controls, there were no significant relationships observed between the FA in the right SLF in kindergarten and subsequent word reading abilities. By contrast, FDR-corrected correlations were observed among at-risk children between FA in the posterior segment of the right SLF and decoding skills (as indicated by the untimed Word Attack subtest from the *WRMT-III; r* = 0.49, *p* = 0.005). These findings are displayed in Figure 1.

### Which kindergarten-age factors best predict subsequent word reading abilities among at-risk children?

In a final step, multiple regression analysis was conducted to evaluate which behavioral and/or neural variables at the kindergarten age best predict decoding abilities among at-risk children only. Based on findings from the correlation analyses, the Word Attack subtest from the *WRMT-III* was selected as the dependent variable. Predictors of interest included factors identified to differ between at-risk children with versus without subsequent dyslexia in the MANOVAs (nonverbal cognitive abilities, SES, and speech production accuracy), as well as key predictors of subsequent literacy skills as previously reported in the literature (phonological awareness, rapid automatized naming, letter-sound knowledge). Control predictors, gender and age at first time point, were also included in the model. For parsimony, a final model was performed without the inclusion nonverbal cognitive abilities, as this predictor did not demonstrate a significant association with the outcome variable in the provisional model. The final multiple regression model explained 80.3% of the variance in decoding abilities at the end of second grade (see Figure 2). This model revealed that the following predictors significantly explained decoding abilities: the posterior segment of the right SLF, age, gender, SES, and phonological awareness (*F* = 10.73, *p* < 0.0001; all individual associations outlined in Table 3).

**Figure 2 Legend:**
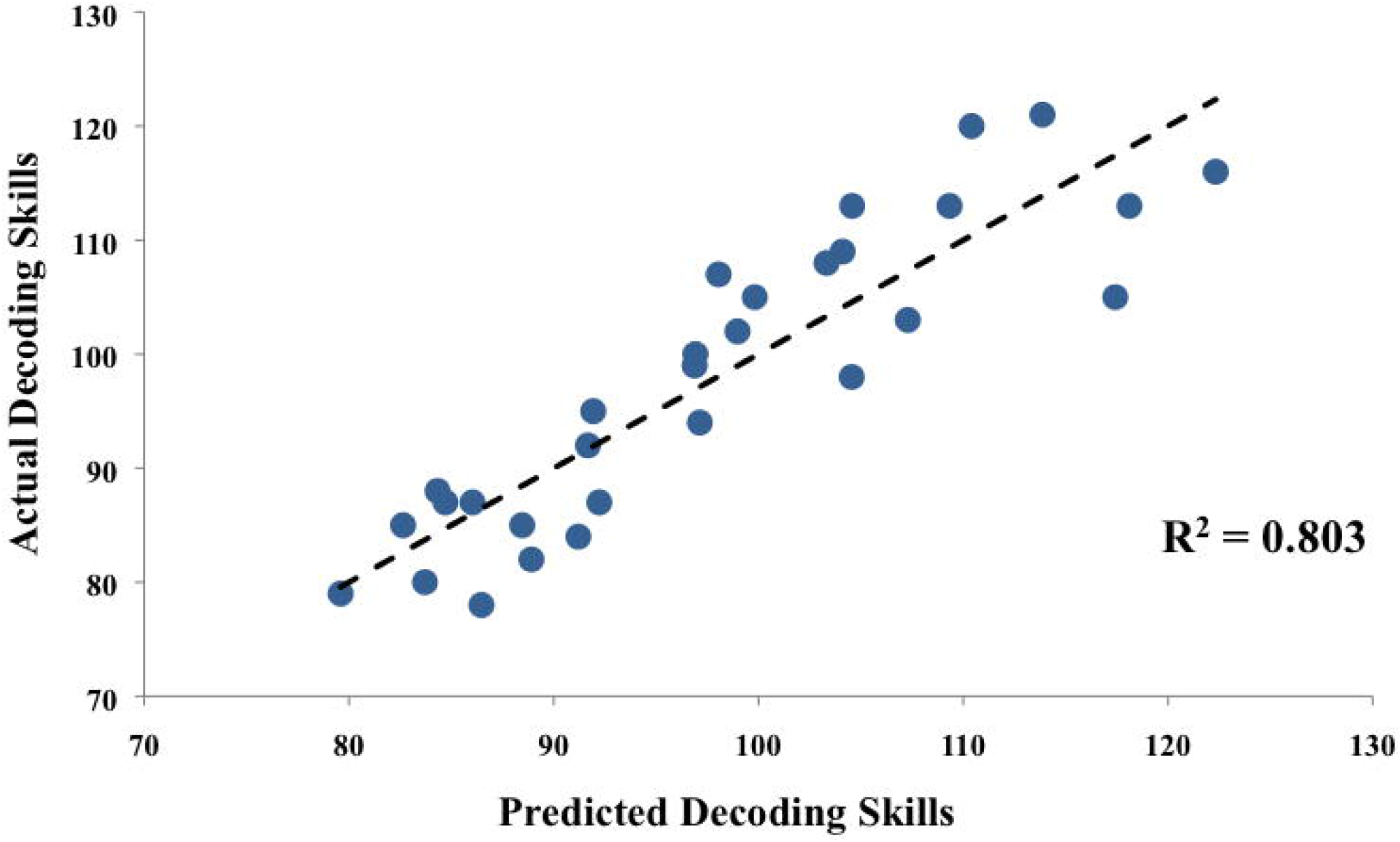
Prediction of decoding skills (Word Attack subtest from the *WRMT-III*) at the end of second grade among at-risk children utilizing multiple regression analysis. Predicted scores are shown on the x-axis, and actual (measured) scores are shown on the y-axis.

**Table 3.** Multiple Regression: Longitudinal Prediction of Decoding Abilities (end of second grade)

| | *B* | *SE B* | $\beta$ | *t* | *p* |
| --- | --- | --- | --- | --- | --- |
| *Putative Protective Predictors* |  |  |  |  |  |
| Right SLF | 73.62 | 21.97 | 0.37 | 3.35 | 0.003 |
| Socioeconomic Status | 0.45 | 0.15 | 0.43 | 3.03 | 0.006 |
| Speech Accuracy | 0.41 | 0.22 | 0.23 | 1.84 | 0.080 |
| *Control Predictors* |  |  |  |  |  |
| Age | -1.03 | 0.38 | -0.33 | -2.74 | 0.010 |
| Gender | 15.86 | 2.91 | 0.62 | 5.44 | <0.001 |
| *Key Pre-Literacy Predictors* |  |  |  |  |  |
| Phonological Awareness | 2.36 | 1.01 | 0.29 | 2.34 | 0.029 |
| Rapid Automatized Naming | 0.05 | 0.15 | 0.04 | 0.33 | 0.742 |
| Letter-Sound Knowledge | -0.15 | 0.18 | -0.10 | -0.80 | 0.435 |
| Intercept | 57.43 | 39.97 |  | 1.44 | 0.165 |

Note: $R^2 = .803$

## DISCUSSION

Among at-risk children identified by early behavioral screening, this longitudinal investigation revealed several factors on cognitive-linguistic, environmental, and neural levels that were associated with subsequently better word reading outcomes at the end of second grade. As hypothesized, group comparison between at-risk children based on subsequent reading outcomes revealed that at-risk children who did *not* develop dyslexia exhibited significantly higher socioeconomic status, speech production accuracy, and microstructure in the right superior longitudinal fasciculus in kindergarten. In addition, the two at-risk groups did not significantly differ on language measures at the start of reading instruction, but differences in vocabulary knowledge emerged by the end of second grade. Moreover, among all at-risk children (regardless of whether they developed dyslexia or not), positive relationships were observed between the right superior longitudinal fasciculus in kindergarten and subsequent non-word reading abilities. This relationship was observed independent of age, nonverbal cognitive abilities, and socioeconomic status, and was not present among typical controls. Lastly, multiple regression revealed that the right superior longitudinal fasciculus, socioeconomic status, phonological awareness, gender, and age in kindergarten significantly predicted subsequent decoding abilities among at-risk children. These findings suggest cognitive-linguistic, environmental, and neural factors already present at the start of formal reading instruction may play a role in promoting reading acquisition among at-risk children.

### Cognitive-linguistic factors associated with better reading outcomes among at-risk children

The cognitive-linguistic factors that differed between at-risk children with versus without subsequent dyslexia in the present study align closely with the prior literature. Speech production accuracy has previously been investigated as a factor that may promote reading acquisition, but the currently available standard articulation tests have only revealed trends towards significant differences between at-risk children with versus without subsequent dyslexia^31,32,35^. In line with previous hypotheses that standard tests may not be sensitive enough to capture group differences in speech production accuracy^31,32^, the present findings identified significant differences through an alternate approach that involved detailed, retrospective analysis of audio-recorded speech samples. Regarding the advantageous role of speech production accuracy in reading acquisition, a relative strength in this domain could facilitate phonological awareness development early on, as preschool-age children with speech delays have shown concurrent improvements in speech accuracy and phonological awareness in intervention that targets both skills^62^. Moreover, phonological awareness has been shown to mediate the relationship between deficits in speech production and subsequent reading outcomes^63^. Although better speech production accuracy was observed among at-risk children with higher reading achievement in the present findings, it is important to note that speech accuracy did not significantly contribute to the prediction of subsequent decoding abilities in the multiple regression analysis. Therefore, the present study contributes to the growing body of evidence suggesting that better speech production accuracy among at-risk children is associated with better reading outcomes, but the direct role that speech accuracy may play in shaping subsequent reading abilities remains unclear.

Although sentence comprehension and vocabulary knowledge did not differ at the start of formal reading instruction between at-risk children who subsequently developed dyslexia compared to those who did not, significant differences in vocabulary did emerge by the end of second grade. The emergence of this difference over the course of learning to read may point towards a consequence of the difficulty with learning to read, as there is a documented gap in reading exposure between those with and without reading deficits^64^. Furthermore, children’s reading abilities have been shown to determine how much they choose to spend time reading^65^. This gap in the time spent reading can result in a cascading effect known as the Matthew effect in reading, in which good readers gain an advantage over poor readers by reading and learning from text^66^. Consequently, children with diminished reading exposure due to the associated difficulties may miss opportunities to acquire new vocabulary once they reach the academic stage of relying on reading to learn new content^67^, which is expected by the end of second grade^65,68^. In line with this notion, vocabulary growth has been shown to be a more important predictor of subsequent language and reading development than initial vocabulary size at early stages in development^69,70^. These findings also support the notion that at-risk children with relative strengths in vocabulary knowledge may be able to discern context clues in text to achieve word identification, despite decoding difficulties^33^. The emerging differences in vocabulary observed over the course of reading acquisition further illuminate the importance of early identification, and suggest that incorporating vocabulary targets in intervention may also be necessary in order to close the gap in exposure that can occur during reading development.

### Socioeconomic status: an important environmental factor linked with reading outcomes

Another undeniably important factor identified in the present study is socioeconomic status (SES), since significantly higher SES was observed among at-risk children who did not develop dyslexia when compared to those who did and SES was found to significantly predict subsequent decoding outcomes among at-risk children. These findings align with the previously reported crucial impact of socioeconomic factors on language and reading development^38–41,71–73^. The true nature of this effect is difficult to interpret, as SES is deeply intertwined with several aspects of the home environment and access to educational resources. This may reflect the differences in reading achievement that can arise due to qualitative aspects of the home and school environment, which include parent support, tutoring options, and even access to books, etc., to name a few^74^. Interestingly, the early impact of SES was not pervasive in the present sample since there were no significant differences in vocabulary knowledge at the start of formal reading instruction, a factor that has previously been shown to distinguish children from high versus low socioeconomic backgrounds early on^43,44^. That said; the emergence of differences in language and reading abilities by the end of second grade aligns with the well-established achievement gap characterized by disparities in SES^45–47^. However, this is also a reflection of the differences in achievement that can emerge as a direct consequence of difficulty with reading^66^. Ultimately, the present findings illuminate the importance of socioeconomic factors among at-risk children in particular, as socioeconomic status at the start of formal reading instruction did distinguish those who subsequently developed dyslexia from those who did not. This finding underscores the importance of characterizing SES in determining approaches to early intervention among at-risk children.

### Right-hemispheric white matter microstructure at the start of formal reading instruction is linked with subsequent reading outcomes among at-risk children

In addition to cognitive-linguistic and environmental factors identified, the present study also observed the previously proposed alterations in underlying neural pathways in right-hemispheric white matter, specifically in the posterior segment of the superior longitudinal fasciculus (SLF). This pathway was first suggested as a potential compensatory neural mechanism, as higher indices of fractional anisotropy (FA) in the right SLF among middle school-age children with dyslexia were positively linked with subsequent word reading abilities approximately two years later^57^. In line with this, children with dyslexia who received intervention and demonstrated significant improvements in word reading have shown increased activation in right-hemispheric regions^75–78^. These activation changes were specific to children who responded to the treatment and were not observed among those who did not improve after remediation, suggesting that increased right-hemispheric activation may serve as a compensatory neural mechanism to facilitate reading development^79^. A longitudinal study starting at the pre-reading stage observed that temporoparietal segments of the right SLF showed a higher rate of development (i.e., increasing FA) over the course of learning to read among children with familial risk for dyslexia who subsequently became good readers relative to those who became poor readers^59^. Building on this evidence, the present study is the first to identify that FA in the posterior (temporoparietal) segment of the right SLF *at the start* of formal reading instruction significantly differs between at-risk children who do *not* develop dyslexia and those who do subsequently develop dyslexia, and significantly predicts subsequent decoding abilities among at-risk children.

At-risk children without subsequent dyslexia showed significantly higher FA in this tract relative to typical controls as well. A positive relationship between microstructure in the right SLF and subsequent non-word reading (decoding) abilities at the end of second grade was specific to at-risk children *only*, as there was no relationship observed among typical controls. These findings are in line with previous work^51,57,59^, and our results demonstrate that it is not only the rate of development in the right SLF over the course of learning to read, but that higher FA in this tract at the time a child begins formal reading instruction is significantly related to better word reading outcomes later on. The specificity of relationships between the right SLF and subsequent decoding abilities among at-risk children only suggests that at-risk children who acquire better word reading abilities may be utilizing the right-hemispheric correlate of the left-hemispheric ‘indirect route’ for decoding that has been previously characterized in typical reading development^80^. Furthermore, this significant relationship was independent of SES, which suggests that the trajectory of at-risk children in the present study is not solely explainable by socioeconomic factors.

### Findings point towards specific protective or compensatory factors that facilitate reading development

Although this work illuminates several factors that seem to be linked with subsequently better word reading outcomes among at-risk children, a critical question remains: were the children identified as at-risk who did not subsequently develop dyslexia ever *truly* at risk? Although major efforts are underway to optimize the accuracy of the early identification of at-risk children at the start of formal reading instruction, current approaches are limited by poor specificity rates, which lead to the over-identification of at-risk children^25–27^. Therefore, it remains unclear whether these children were the result of false-positive identification, and would have learned how to read without difficulty regardless of early identification. Alternatively, it is possible that certain protective or compensatory strategies may ‘override’ current approaches to early screening by promoting reading acquisition among at-risk children. After all, the present findings identified multiple factors at cognitive-linguistic, neural, and environmental levels that distinguished at-risk children based on their subsequent reading outcomes, which is in line with the hypothesis that certain risk factors increase the liability for dyslexia, whereas putative ‘protective’ factors may decrease the liability^28,29^. In addition, at-risk children without subsequent dyslexia showed significantly higher FA in the right SLF than typical controls, which suggests that this group does not likely encompass only false-positive identifications, but points towards a protective or compensatory neural mechanism among at least some at-risk children who did not develop dyslexia. Moreover, the positive relationships between FA in the right SLF in kindergarten and subsequent decoding abilities among at-risk children *only* suggests that the higher the FA in this tract, the better at-risk children have done in acquiring decoding abilities over time.

An important ongoing challenge is to determine the specific developmental timeframe in which early screening may be most reliable and effective. Recognizing the rapid development of the behavioral skills that are assessed in early screening at the preschool/kindergarten age^81^, screening conducted at the emergent reading stage based on word reading abilities has shown accurate prediction of subsequent outcomes^82^. Accordingly, multi-stage screening over the course of reading acquisition has been proposed when feasible, which has been shown to minimize the rate of false-positive identifications^82,83^. Yet, additional evidence suggests that intervention is more effective when provided earlier rather than later^24,84–87^. Moreover, the present findings of a gap in language achievement over the course of learning to read further point towards the importance of early screening in order to prevent this effect and potentially facilitate compensatory strategies. While reducing the prevalence of over-identification in early screening is necessary for optimal allocation of resources, it seems especially important to ensure that children who may benefit from early intervention are provided with this opportunity in order to promote better outcomes.

### Future Directions

The present study lays important groundwork for several future directions. Cognitive-linguistic, environmental, and neural factors evident at the start of formal reading instruction seem to be linked with subsequent word reading outcomes among these at-risk children, however, it remains unclear how early in development these may have arisen. The field has largely considered these ‘protective’ factors, yet we do not know whether factors are evident from infancy that truly ‘protect’ at-risk children throughout subsequent development, or whether certain compensatory strategies/experiences that arise throughout early childhood shape the developmental trajectory of reading acquisition^51^. More longitudinal studies starting in infancy are needed to establish this neurodevelopmental trajectory and determine the onset of these putative factors. For instance, direct comparison of infants with and without familial risk for dyslexia in this context will be necessary to determine the potential etiologic basis of the right-hemispheric pathway identified. In addition, there are several potential contributing factors that were not addressed in the present analysis. It is possible that certain aspects of the home environment may have played an important role (particularly pertaining to literacy exposure and enrichment), or access to tutoring and early intervention/targeted instruction. No differences in the amount of tutoring or early intervention received by at-risk children in the present study were observed, however, qualitative aspects have yet to be investigated. While these factors warrant further investigation in future work, the focus of the present study makes an important contribution by identifying that multidimensional factors, present when at-risk children start receiving reading instruction, are linked with their subsequent word reading outcomes.

### Clinical Implications and Conclusions

Overall, this work carries several important implications for approaches to early identification and intervention to promote reading acquisition among at-risk children from the start of formal reading instruction. The present findings point towards several specific strategies that may be employed to best serve at-risk children identified based on current methods of early screening, such that if a child is identified as at-risk based on the key predictors of dyslexia (phonological awareness, rapid naming, letter identification, oral language skills), then next steps would involve the following: (a) refer this child to evaluation by a speech-language pathologist to consider relative strengths or weaknesses in the areas of speech and language, (b) assess nonverbal cognitive abilities to determine whether this may be a relative strength or weakness, and (c) acquire information about the home environment in order to identify whether this will likely be a source of support, or to consider approaches to supporting the home environment through educational and clinical services provided to parents. From there, continued progress monitoring and implementation of multi-stage screening are recommended whenever feasible, to minimize inaccurate allocation of services to those who may not need it^82,83^. Ultimately, this approach offers great promise to promote optimal outcomes when children are then provided with early intervention that targets each child’s specific risk profile *and* builds upon areas of strength as potential protective factors or compensatory strategies.

For at-risk children who do receive targeted instruction or clinical services thereafter, the present findings support the notion that multiple approaches to targeted instruction in the first few years of formal schooling may be particularly effective. In addition to the well-established goal to promote decoding skills, there is potential to reach another level of support through careful consideration of what may be most effective for each child based on a holistic profile of their relative strengths and weaknesses. Ultimately, this study provides a better understanding of factors on cognitive-linguistic, environmental, and neural levels that may promote better word reading outcomes for at-risk children from the start of formal reading instruction.

## METHODS

### Participants and design

Seventy-four children (37 female / 37 male, age range: 4:9 – 6:6 [yrs: mo], mean age: 5:6, STD: 4.12 months) were included in the present study as part of a larger longitudinal investigation of children at behavioral risk for dyslexia Researching Early Attributes of Dyslexia (i.e., the READ study)^88-91^. Initially, screening for early risk for dyslexia was conducted in pre-kindergarten and kindergarten classrooms in New England, from schools which were diverse in type (public district, public charter, private, and religious) and location (urban and suburban areas). Screening took place in the spring of prekindergarten or early fall of the kindergarten year, before or immediately after the start of formal reading instruction. Consequently, a subset of these children were enrolled in the longitudinal aspect of this study (*n* = 186 in total), which involved further behavioral assessment as well as neuroimaging in kindergarten, followed by behavioral re-evaluation until the end of second grade. Children identified as at-risk were oversampled in the selection of longitudinal enrollment to ensure variance in subsequent reading outcomes. Children included in the present study were right-handed, native American English speakers with no history of neurological or psychiatric disorders, head or brain injury, or poor vision/hearing per parent report. In addition, only children who demonstrated nonverbal cognitive abilities with a standard score of at least 80 (as indicated by the *Kaufman Brief Intelligence Test: 2nd Edition* KBIT-2^92^ were included in the present study. Moreover, the present study focused solely on children in the sample for whom quality diffusion imaging data was acquired who also had completed longitudinal follow-up investigation.

Parents completed questionnaires detailing the child’s handedness, family history of language and/or learning disabilities, and socioeconomic status (as indicated by the Barratt Simplified Measure of Social Status^93^, which derives a composite score for mother’s and father’s education levels and occupation). Informed consent was obtained by all participants, such that verbal assent was obtained from each child and written consent from each legal guardian, respectively. All experimental protocols and procedures were approved by The Institutional Review Boards at Boston Children’s Hospital and Massachusetts Institute of Technology, and all research was performed in accordance with relevant guidelines and regulations.

### At-risk classification based on early screening

Initial screening took place before or within the first few weeks of formal reading instruction (e.g., pre-kindergarten and kindergarten in the United States), utilizing the primary behavioral predictors of dyslexia to classify risk status. These included measures of phonological awareness, rapid automatized naming, and letter identification. Children who demonstrated performance below the 25 ^th^ percentile on any of these three classification measures within the large sample initially screened (*n* = 1,433) were categorized as ‘at-risk.’ Children who demonstrated performance above the 25^th^ percentile on these measures were not identified as at-risk.

### Classification measures

Phonological awareness was measured by the composite of the Elision, Blending Words, and Nonword Repetition subtests from *The Comprehensive Test of Phonological Processing* CTOPP^94^. Elision measures the ability to segment and delete phonemes from a word, and Blending requires children to combine a string of individually presented speech sounds to form a word. Nonword Repetition has children repeat recorded nonwords as accurately as possible. The mean of these three subtests was used to calculate a phonological awareness composite score. Rapid automatized naming was measured by the Objects and Colors subtests from the *Rapid Automatized Naming/Rapid Alternating Stimulus* (RAN/RAS) tests^95^, which measures accurate, rapid naming of familiar items in an array. For children who demonstrated knowledge of letters, the Letters subtest was also acquired for RAN/RAS tests. The mean of all subtests administered comprised the Rapid Automatized Naming composite. Letter knowledge was characterized by scores on two subtests: the Letter Identification subtest from the *Woodcock Reading Mastery Tests-Revised, Normative Update* (WRMT-R NU^96^), which measures the ability to name letters of the English alphabet. In addition, the Letter Sound Knowledge subtest from the York Assessment of Reading for Comprehension (YARC^97^) was administered, which assesses knowledge of letter sounds. Note that the YARC was normed based on a UK sample, which did not seem representative of our US-based sample. Therefore, norms for this measure were established based on the sample distribution of the present sample^89^. The mean of standard scores for these two subtests derived the letter knowledge composite score.

Additional constructs that may impact a child’s risk status were evaluated at initial screening. Specifically, syntactic language abilities were screened utilizing the Sentence Repetition subtest of the *Grammar and Phonology Screening* (GAPS^98^), in which performance below the 10^th^ percentile on this measure indicated significant syntactic language difficulties. In this subtest, children are asked to directly imitate sentences presented orally. Word reading ability was also characterized to account for potential word reading competency at the time of screening. This was measured with the Word Identification subtest from the WRMT-R NU, which assesses single real word reading skills in an untimed manner.

### Behavioral measures of interest

#### Additional assessment in kindergarten

Children enrolled in the longitudinal aspect of the study who underwent neuroimaging also returned in kindergarten to complete a more comprehensive battery of standardized assessments. A subset of these assessments were selected based on the focus of the present study, which included measures of speech and language hypothesized to serve protective/compensatory roles based on the previous literature. Measures included were as follows:

##### Components of language

Standardized measures were utilized to evaluate two aspects of language. Vocabulary knowledge was characterized by *The Peabody Picture Vocabulary Test* (PPVT-4^99^), in which children indicated which of four pictures best represents the word said by the examiner. Sentence comprehension was characterized by the sentence structure subtest from the *Clinical Evaluation of Language Fundamentals* (CELF-IV^100^), in which a child is asked to select which of four pictures best represent the sentence said by the examiner.

##### Speech production accuracy

Speech production accuracy was retrospectively determined through audio-recorded connected speech samples from the battery of standardized assessments. Specifically, audio recordings of the GAPS sentence repetition and CTOPP Elision subtests were reviewed and analyzed for phonetic accuracy through Percent Consonants Correct-Revised (PCC-R) analysis by researchers with training in speech-language pathology and linguistics^101^. Per PCC-R procedures, distortions were not counted as errors. In addition, productions characteristic of the New England regional dialect were not counted as errors^102^. Inter-rater reliability was conducted between two raters with 15% of the sample to verify consistent identification across raters, and an intraclass correlation coefficient > 0.9 was achieved. Speech production accuracy was then quantified by the percentage of the number of correct consonants divided by the number of correct and incorrect consonants produced.

#### Longitudinal follow-up at the end of second grade

Follow-up measures acquired at the end of second grade selected for the present investigation focused on subsequent language and word reading abilities. In terms of language, vocabulary knowledge (PPVT-4) and sentence comprehension (sentence structure subtest from CELF-IV) were re-assessed. Word reading abilities were characterized by a series of subtests evaluating word identification and nonword decoding measures in timed and untimed conditions. For timed measures, The Sight Word Efficiency and Phonemic Decoding Efficiency subtests of the *Test of Word Reading Efficiency* (TOWRE-2^103^) were included to measure the ability to read single words rapidly and accurately, in which children read aloud lists of words or non-words, as quickly as possible. Untimed measures included the Word Identification and Word Attack subtests of the *Woodcock Reading Mastery Tests,* Third Edition WRMT-III; ^104^, which required children to read words or non-words aloud that became increasingly more complex.

#### Classification of subsequent word reading abilities

To align with the group comparison approach employed in related previous literature, children were classified based on risk status in kindergarten determined by initial screening and subsequent reading abilities at the end of second grade. Classification was established for the following three groups: at-risk kindergarteners who subsequently developed dyslexia, at-risk kindergarteners who subsequently did *not* develop dyslexia, and typical controls (no risk, typical readers). Classification of subsequent word reading outcomes was determined based on the definition of dyslexia traditionally utilized for diagnosis^105^. Therefore, children met the criteria for dyslexia if they demonstrated performance one standard deviation below the mean or lower (standard score ≤ 85) on at least one subtest of the timed and untimed standardized word reading measures administered at the end of second grade (TOWRE-2 and WRMT-III). Children who had standard scores of 90 and higher on all word-reading measures were classified as typical controls.

Additional potential risk factors were carefully considered to ensure appropriate group classification. In terms of potential co-occurring syntactic language difficulties, screening with the Sentence Repetition subtest of the GAPS revealed that four children in the present sample demonstrated scores below the 10^th^ percentile. However, all of these children were already classified as ‘at-risk’ based on the core behavioral predictors (phonological awareness, rapid naming, letter identification). As for children who were not classified as at-risk in kindergarten but then did go on to develop dyslexia, an insufficient number were identified to warrant inclusion in the present study (*n* = 6). In addition, a subset of children reportedly had family history of dyslexia but otherwise presented as typical controls (e.g., not classified as at-risk based on early screening and subsequently developed typical reading abilities, *n* = 12). Given the reported history of familial risk, these children were not included in the typical control group.

Therefore, the present investigation examined the following three groups: at-risk kindergarteners who subsequently developed dyslexia (*n* = 17), at-risk kindergarteners who subsequently did *not* develop dyslexia (*n* = 18), and typical controls (*n* = 39). These groups were age-matched to rule out the possibility of an age effect in the analyses. With 17 children in the present sample having met the criteria for dyslexia, the prevalence rate for dyslexia is 22% in this sample, which aligns appropriately with the oversampling approach taken in the selection of at-risk children longitudinally enrolled to achieve a representative distribution.

#### Neuroimaging acquisition

Children were first introduced to the MR scanner setting with a child-friendly mock scanner training, which allowed them to acclimate to the MR environment. Structural neuroimaging was acquired as one aspect of a 40-minute imaging protocol, which included breaks as individually requested, on a 3T Siemens Trio Tim MRI scanner with a standard Siemens 32-channel phased array head coil. A structural T1-weighted whole-brain anatomical volume was acquired (multiecho MPRAGE; acquisition parameters: TR = 2350ms, TE = 1.64ms, TI = 1400ms, flip angle = 7°, FOV = 192 × 192, 176 slices, voxel resolution = 1.0 mm3, acceleration = 4). An online prospective motion correction algorithm was employed to mitigate motion artifacts. To further monitor potential motion during acquisition, a researcher stood near each child in the MRI room to present a physical reminder to stay still throughout the session when necessary. Diffusion-weighted images were acquired with 10 non-diffusion-weighted volumes (*b* = 0) and 30 diffusion-weighted volumes (acquisition parameters: *b* = 700 s/mm^2^, 128 × 128 mm base resolution, isotropic voxel resolution = 2.00 mm^3^).

#### Diffusion weighted image processing and automated fiber quantification

Diffusion weighted images were processed with the approach taken in a previous study from the lab with a cohort of children of the same age. A brain mask was generated from the structural T1-weighted image utilizing the Brain Extraction Tool (BET^106^) to separate brain tissue from non-brain tissue. Raw DWI data were converted from DICOM to NRRD format through the DICOM-to-NRRD Converter software of Slicer4 (www.slicer.org). DTIprep software^107^ and visual inspection were used to evaluate scan quality. The translation threshold was set to 2.0 mm and the rotation threshold to 0.5° for determining motion artifacts. Volumes containing motion artifacts were removed prior to diffusion tensor estimation. After assessing scan quality, DWI data were processed with the VISTALab mrDiffusion toolbox and diffusion MRI software suite (www.vistalab.com), including eddy current correction and tensor-fitting estimations with a linear least-squares (LS) fit for fitting the diffusion tensors.

White matter tracts were identified using the Automated Fiber Quantification (AFQ) software package (github.com/jyeatman/AFQ)^108^. For a full description of the procedures and parameters used, see Wang and colleagues (2016). For a brief summary, whole brain tractography was computed using a deterministic streamline tracking algorithm^109,110^. Fractional anisotropy (FA) was then computed for each tract based on eigenvalues from the diffusion tensor estimation^111^. Fiber tracking was conducted with an FA threshold of 0.2, and stopped in cases where the minimum angle between the last path segment and the next step direction was > 40°. Region of interest (ROI)-based fiber tract segmentation and fiber-tract cleaning were then employed (using a statistical outlier rejection algorithm), and then FA quantification was conducted along the trajectory of each fiber. FA of each fiber was sampled to 100 equidistant nodes.

The present study focused solely on the right-hemispheric superior longitudinal fasciculus, to build upon previous studies implicating this tract as a specific protective or compensatory neural mechanism that relates to better word reading outcomes among children at risk and those with dyslexia ^57,59^. For this tract, the characterization of 100 nodes was resampled to 50 nodes by discarding the portion of the tract where individual fibers separated from the core fascicle toward their destination in the cortex. This method was employed based on Wang and colleagues^59^, as it is known to improve normalization and co-registration of each tract for group comparisons^112^. FA values for each node along the trajectory of the tract were then averaged into three segments to maximize statistical power.

### Statistical analyses

#### Group comparisons

To address the main research question, at-risk children who did *not* subsequently develop dyslexia were directly compared to at-risk children who did develop dyslexia as well as typical controls on factors evaluated at cognitive-linguistic, environmental, and neural levels. Direct comparisons between these three groups were evaluated accordingly through a one-way MANOVA with measures of interest at the kindergarten age, and then the end of second grade as well. Significant variables identified in the MANOVAs were then further examined through post-hoc pairwise comparisons, controlling for the False Discovery Rate (FDR) to correct for multiple comparisons^113^.

#### Correlations between the right SLF in kindergarten and subsequent word reading abilities

To further examine relationships between the right superior longitudinal fasciculus (SLF) in kindergarten and subsequent word reading abilities among at-risk children, partial correlations were employed. Partial correlations were conducted to control for the potential effect of the following demographic variables: age, nonverbal cognitive abilities, and socioeconomic status (as indicated by the index of total parent education and occupation on the Barratt Simplified Measure of Social Status). Specifically, partial correlations were conducted between the average FA for each segment of the SLF specified (anterior, mid, posterior) and each subsequent word reading measure acquired at the end of second grade (subtests from the *TOWRE* and *WRMT-III*). Based on the hypothesis that these relationships would be specific to at-risk children, partial correlations were first conducted among at-risk children only. Relationships were then examined among typical controls. FDR correction for multiple comparisons was employed.

#### Multiple Regression Analyses

To further examine which behavioral and/or neural variables at the kindergarten age best predict word-reading outcomes among at-risk children, multiple regression analyses were conducted among at-risk children only. The dependent variable, word reading performance at the end of second grade, was specified based on the results of the partial correlation analyses conducted among at-risk children only. Predictor variables included demographic variables (age, gender), key precursor literacy skills (phonological awareness, rapid automatized naming, letter-sound knowledge), and putative ‘protective’ factors identified to differ between at-risk children with versus without subsequent dyslexia in the MANOVAs. All statistical analyses were performed utilizing IBM SPSS Statistics software (Version 23 ^114^).

## Acknowledgements

We thank all participating families for their longitudinal commitment to this study, and school coordinators and principals who made screening possible (for an overview of participating schools, see http://gablab.mit.edu/index.php/READstudy). We are grateful for all additional members of the READ team who contributed to data collection, especially Sara Beach, Abigail Cyr, and Kelly Halverson. We also wish to acknowledge the Athinoula A. Martinos Imaging Center at the McGovern Institute for Brain Research at MIT. This work was supported by the Eunice Kennedy Shriver National Institute of Child Health and Human Development at the National Institutes of Health (R01 HD067312). Funding was also provided for Jennifer Zuk by the National Institutes of Health National Research Service Award (F31 DC015919-01), the American Speech-Language-Hearing Foundation, and the Sackler Scholar Programme in Psychobiology.

## Data Availability Statement

The datasets generated during and/or analyzed during the current study are available from the corresponding author on reasonable request.

## Author Contributions

N.G., J.G., and E.N. designed research; J.Z., T.H., N.G., and E.N. designed retrospective research; J.Z., J.D., E.N., X.Y., O.O.P. and Y.W. performed research; J.Z., N.G., T.H., J.D., E.N., O.O.P., X.Y., and Y.W. analyzed data; J.Z., J.D., X.Y., N.G., T.H. and J.G. wrote the paper.

## Competing Interests

The author(s) declare no competing interests.

